# *Acinetobacter baumannii*’s Great Escape: How a Membrane Protein Interacts with Fibronectin to Evade Immunity

**DOI:** 10.1101/2024.12.26.630398

**Authors:** Laurine Vasseur, Florent Barbault, Antonio Monari

## Abstract

Bacterial resistance and nosocomial infections are serious threats compromising public health in numerous countries. Acinetobacter baumannii has been identified as one of the most serious pathogens, due to its potential virulence and the development of multiple resistance. In this contribution by using all-atom molecular dynamics simulation, we analyze the structure and the properties of the OmpA protein, which is present in the bacterial external membrane. We also analyze the structure of possible protein/protein complexes formed between OmpA and the human fibronectin, which may ultimately lead to the immune escaping of the bacteria. For the first time, we provide for the first time plausible structure of the complex also identifying suitable amino acid mediating the interaction, and thus, constituting suitable drug targets to disrupt the complex formation.

**TOC Abstract:** 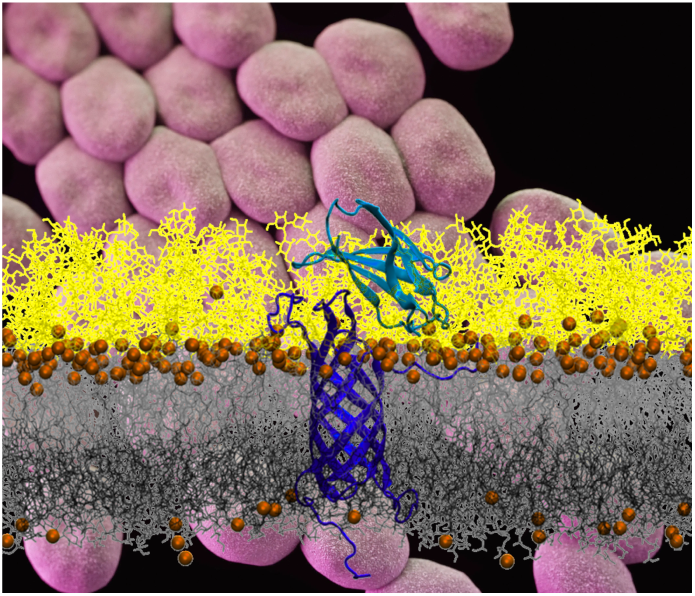

## INTRODUCTION

*Acinetobacter baumannii* (*A. baumannii*) is a Gram-negative bacterium which is raising significant concern worldwide mainly due to its capacity to induce nosocomial infections^1–3^. Diseases resulting from the exposition to *A. baumannii* include pneumonia, urinary tract infections, meningitis, and septicemia^4,5^. Furthermore, such infections may be particularly severe and correlate to very bad prognosis due to the bacterial high resistance to antibiotics which leads to high mortality rates^6^. Hospitalized patients, particularly in intensive care units (ICUs), are exposed to the highest risk, as the bacteria invade the organisms penetrating through eventual skin lesions and the respiratory tract^4,7^. As a matter of fact, ventilator associated pneumonia (VAP), which is favored by mechanical ventilation, is the most common nosocomial infection caused by *A. baumannii*, with mortality rates ranging from 40% to 70%^8,9^. Moreover, the spread of *A. baumannii*’s strains showing antibiotic multi resistance has emerged as a most stringent global concern. Indeed, the World Health Organization (WHO) has classified *A. baumannii* as a member of the ESKAPE group of microorganisms (*Enterococcus faecium, Staphylococcus aureus, Klebsiella pneumoniae, A. baumannii, Pseudomonas aeruginosa and Enterobacter spp*.), which represent major therapeutic threats^10,11^. Consequently, the development of novel antibiotic or therapeutic strategies counteracting the proliferation and spread of *A. baumannii* is considered as a top priority in the domain of public health.

Differently from their Gam-positive counterparts, Gram-negative bacteria possess an envelope constituted by an internal and an external membrane, which define the periplasmatic space in between them. Indeed, the outer membrane, which embeds lipopolysaccharides specific to the bacterial species, forms a complex barrier, which is essential the adaptation to the external environments. Namely, the outer membrane regulates the entry and exit of organic compounds by means of specific proteins, called outer membrane proteins (OMPs).^12,13^ Among them, the OmpA family is a group of surface proteins, which are highly conserved and abundant in Gram-negative bacteria^13,14^. Usually, OmpA proteins act like porin, allowing the outer membrane permeabilization via the formation of water channel reaching the periplasm, and thus the communication with the external medium. In Acinetobacter baumannii, OmpA plays a key role in contributing to antibiotic resistance via the modulation of the membrane permeabilization, and particularly favoring the passive externalization of antibiotics prior to their entrance into the bacterial cell.^14–17^

However, the role of OmpA is wider and more general making this protein essential for bacterial survival. Indeed, it also favors bacterial adhesion, notably in the formation of biofilms, or in promoting cell invasion. Indeed, it has been shown the OmpA interacts with human proteins, notably fibronectin (FN), allowing its adhesion to the host cells^18–20^, but also equipping the bacteria with shields which prevent their recognition by the immune systems^16^, and, thus, favors the spreading of the infection.

FN itself is a protein possessing a fibrillar structure, which is localized in the extracellular matrix (ECM), and plays a central role in regulation of cell adhesion, migration, growth, and differentiation. Thanks to the presence of specific domains, it may bind to various biological macromolecules, such as collagen, integrin and other ECM components^21–23^. Furthermore, several pathogenic bacteria, exploit FN as a receptor and binding spots to firmly adhere to host cells, a crucial step in their invasion and immune evasion^24^. These FN-mediated mechanisms constitute a major virulence factor for many Gram-negative bacteria, enabling them to establish persistent and resistant infections^18,20,25,26^.

It has been shown that in *A. baumannii*, OmpA associates directly with FN, leading to the reinforcement of the bacterial adhesion via further interactions with ECM proteins such as integrins^27–30^, the adhesion of OmpA and FN also correlates with a reduced immune recognition and response. The controlled disruption of the OmpA/FN complex formation, could, thus, be regarded as a most valuable therapeutic strategy to counteract the virulence of *A. baumannii’s* infections. However, the molecular bases by which the two proteins interact and recognize themselves, in the complex and highly inhomogeneous environment of the outer membrane are presently lacking. No structure of OmpA or of the OmpA/FN complex are available to the best of our knowledge. In this contribution we aim to leverage the capacities of long-range molecular dynamics (MD) simulations to achieve μs-scale sampling of the conformational landscape of the two proteins and their complex. This will allow understanding and pointing out the main structural features of the protein/protein interface and the key amino acids driving the complex formations. Therefore, it will also allow to highlight potential druggable regions of the bacterial protein which could lead to the inhibition of the OmpA/FN association and thus may reinstate a correct immune response in case of bacterial infection.

## COMPUTATIONAL METHODOLOGY

All-atom MD simulations have been performed on OmpA embedded in a lipid bilayer mimicking the composition of the *A. baumannii* outer membrane, notably involving two dissymmetric leaflet : the external one composed of *A. baumannii* lipopolysaccharides, and the internal one composed of 74% PVPG, 21% PPPE, and 5% PVCL2.^31^ The initial structure of the bacterial protein has been modeled using Alpha-fold 2.0^2^. OmpA involves a disordered periplasmatic tail which is not correctly reproduced by AlphaFold.^32^ However, since we are mostly interested in the recognition of FN by the outer OmpA domain, the disordered tail has been removed in the following. The initial protein has been embedded in the membrane environment using the Charmm-gui web server^33^. A water buffer has been added including physiological 0.15 M concentration of KCl. Isolated human FN has also been modeled by MD simulation; the initial structure of the protein has been retrieved from the pdb database (1FNF).^34^ Once stable structures of both proteins have been obtained Haddock web server^3^ has been used to make protein/protein docking. It has been performed to identify the most probable interactions pathways between FN and the outer regions of OmpA, Charmm force field has been used to model the proteins, while charm lipid describes the complex membranes and Water modeled with the TIP3P model.^35^ All the MD simulations have been performed using the NAMD 3.0^36,37^ in different independent replicas (see Supplementary Information’s) the results have been analyzed and visualized using VMD^38^ and cpptraj.^39^

## RESULTS AND DISCUSSION

FN is a large weight protein largely present in the extracellular matrix, where it mediates interactions with different membrane receptor, especially integrins. However, FN may also bind to other protein components, including collagen. Structurally, FN is a dimer composed of two long fibrillar, quasi-linear, units bridged together by disulfide bonds. Each unit may be decomposed in globular, beads-like, subunits presenting a high density of β-sheets, tethered by very short, more flexible linkers. A slight variability exists between the different subunits, leading to the classification of type I, II, and III modules. Notably, type III subunits do not present any intrachain sulfur bridges, differently from the other two counterparts. A further variability may also be observed in Type III units due to the presence, or absence of a RGD motif. The RGD motif has been identified as playing an essential role in cellular adhesion, as it is a sequence of three amino acids specifically recognized by a large number of integrins.^40^ These transmembrane proteins are critically involved in cellular adhesion, migration, and signaling.^41^ However, the RGD motif of fibronectin has been shown to be essential for development but not required for fibril assembly.^42^ Based on these information, FN units containing an RGD motif were distinguished from those without to further explore their specific biological roles. Lehay et al. have reported the crystal structure of four FN Type III subunits.^34^ In Figure 1, we report the results of the MD simulations for this simplified model. In particular, we may observe that the high rigidity of the beads is confirmed. Indeed, as shown in Figure 1A their behavior can be approximated as the fluctuations of 4 rigid bodies bridged together by flexible links. This can also be appreciated by the analysis of the Root Mean Square Fluctuation (RMSF) displayed on Figure 1B. More importantly, it also appears that the behavior of the four units is uncoupled and that the FN maintains a globally linear structure with only limited deviations. Obviously, the observed fluctuations are more marked at the terminal ends, testifying to increased flexibility in these regions. As a matter of fact, the same pattern described here is observed also for the other replica, further confirming the rigidity of the structure. These results are not unexpected and is coherent with FN biological role, since it can favor the establishment of a structuring network maintaining cell adhesion. On the other hand, the linear structure of FN, its uncoupled movement, and the structural similarity between the beads justify to simplify the study of the interaction with the bacterial protein, taking int account only one unit at the time.

**Figure 1.**
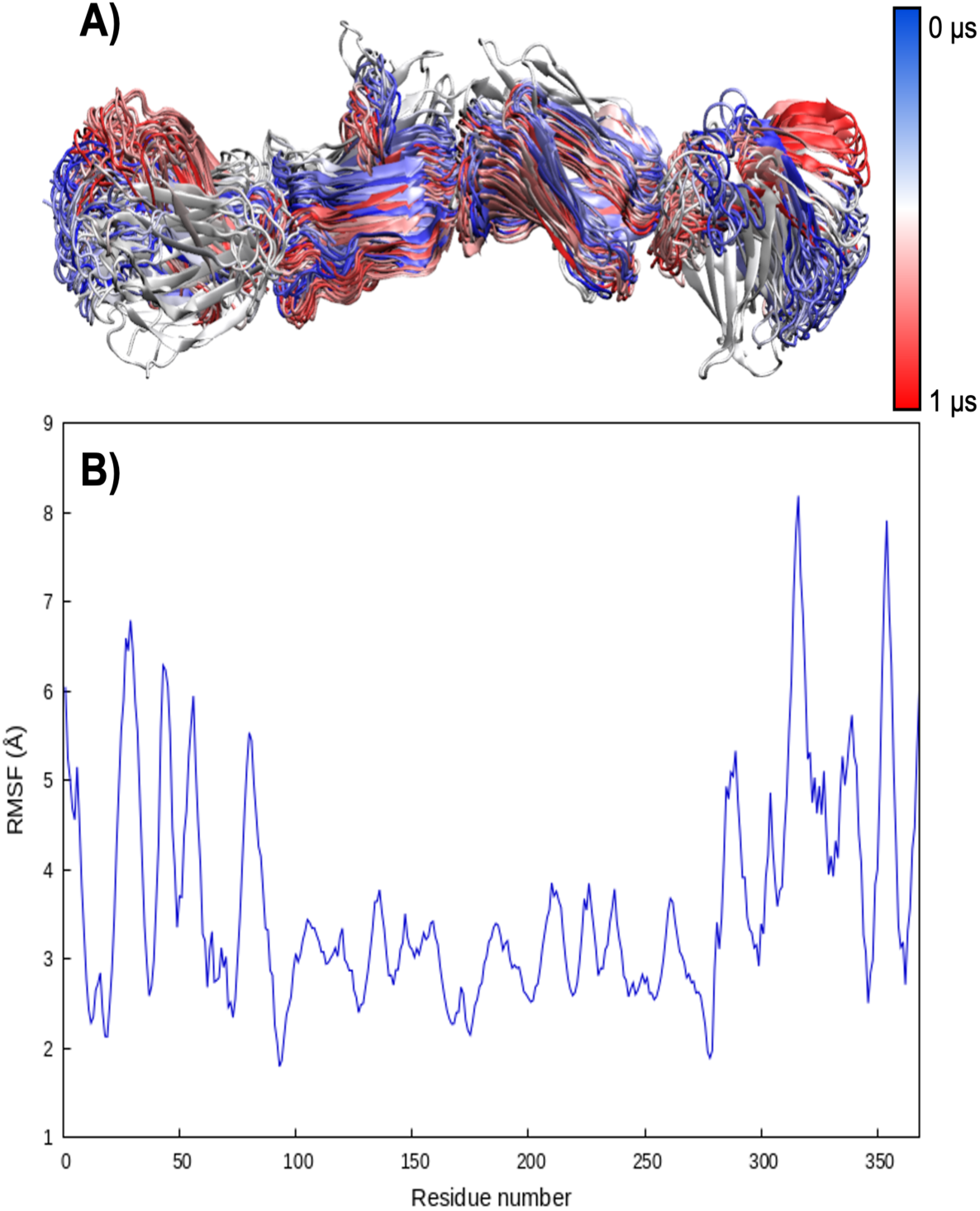
A) superposition of FN structures obtained during the MD simulations for replica 1. B) RMSF of FN averaged on the four replicas calculated for the last 200ns.

The N-terminal transmembrane domain of OmpA has a rather rigid β-barrel structure, flanked by three more flexible arms extending in the polar head region of the outer leaflet. On the other hand, the disordered C-terminal domain extending into the periplasme has been removed from the simulations. The rigidity and the stability of the protein can also be estimated by the analysis of the Root Mean Square Deviation (RMSD, Figure 2A) and RMSF (2B). Note that the sharp increase of the RMSF observed at positions 160 may be assigned to the beginning of the periplasmatic disordered region. The positions of the three arms can also be evidenced by the smaller peaks in the RMSF distribution. These structural features are similar to other bacterial membrane proteins, mediating adhesion and shielding from the immune system, such as in the case of Ail in *Yersinia pestis*.^31^ However, as shown in Figure 2C, the OmpA cavity is sufficiently large and hydrophilic to allow the formation of a continuous water channel, ensuring communication between the periplasm and the extracellular medium. Even if this aspect will not be treated further in the present contribution, it is important to underline that, as a porin, OmpA may participate to the insurgence of antibiotic resistance by favoring the expulsion of potential drugs.

**Figure 2.**
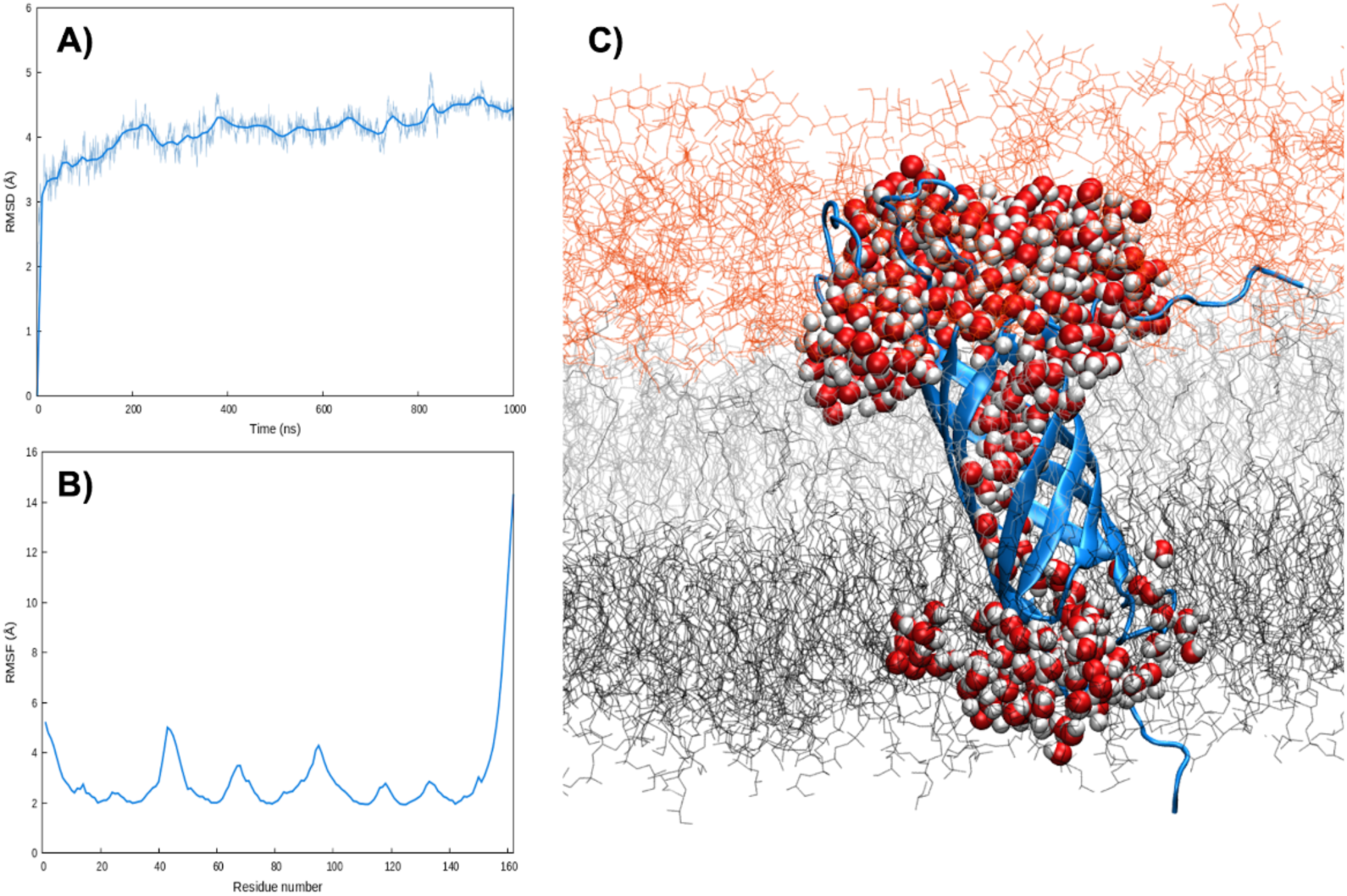
A) Averaged RMSD and B) RMSF of OmpA embedded in the external membrane after equilibration and thermalization. RMSF has been calculated for the last 200ns. C) image of OmpA embedded in the bacterial membrane, a water channel allowing the transmembrane communication is also shown.

To properly characterize the formation of FN/OmpA complex and its stability we firstly performed protein/protein docking. As already said our model involved only one FN subunit. Due to the sequence difference, we performed docking for both the RGD-containing and the non-RGD subunits. As shown in SI two poses stand out for the RGD-containing model, while only one independent conformation can be highlighted for the non-RGD system. These initial structures were engaged in MD simulations.

The results of the MD simulations are collected in Figure 3 and confirms that the complex between the two proteins is stable, for both the RGD-containing and non-RGD unit. The stability of the complex can also be appreciated from the time series of the RMSD for the complex, reported in Figure S2, which shows only very limited oscillations. Interestingly, as can be seen from Figure 3A and 3B, complexes with different orientations of FN are possible, whereas the RGD domain does not appear to play a pivotal role in favoring the interaction. The latter was expected, since OmpA has no structural similarity to an integrin. However, FN is consistently interacting with the flexible arms of OmpA and is positioned in the polar head region (Figure 3D). Not unexpectedly FN clearly embeds in the lipopolysaccharide shield encompassing the external membrane. Note that the same behavior is observed for almost all the replica except for one case, as shown in SI, in which the FN units destabilize the membrane. This corresponds to the conformation that leads to the RGD_1 pose and is probably due to the fact that FN only interacts with one of the flexible arms, thus lacking a firmer support.

**Figure 3.**
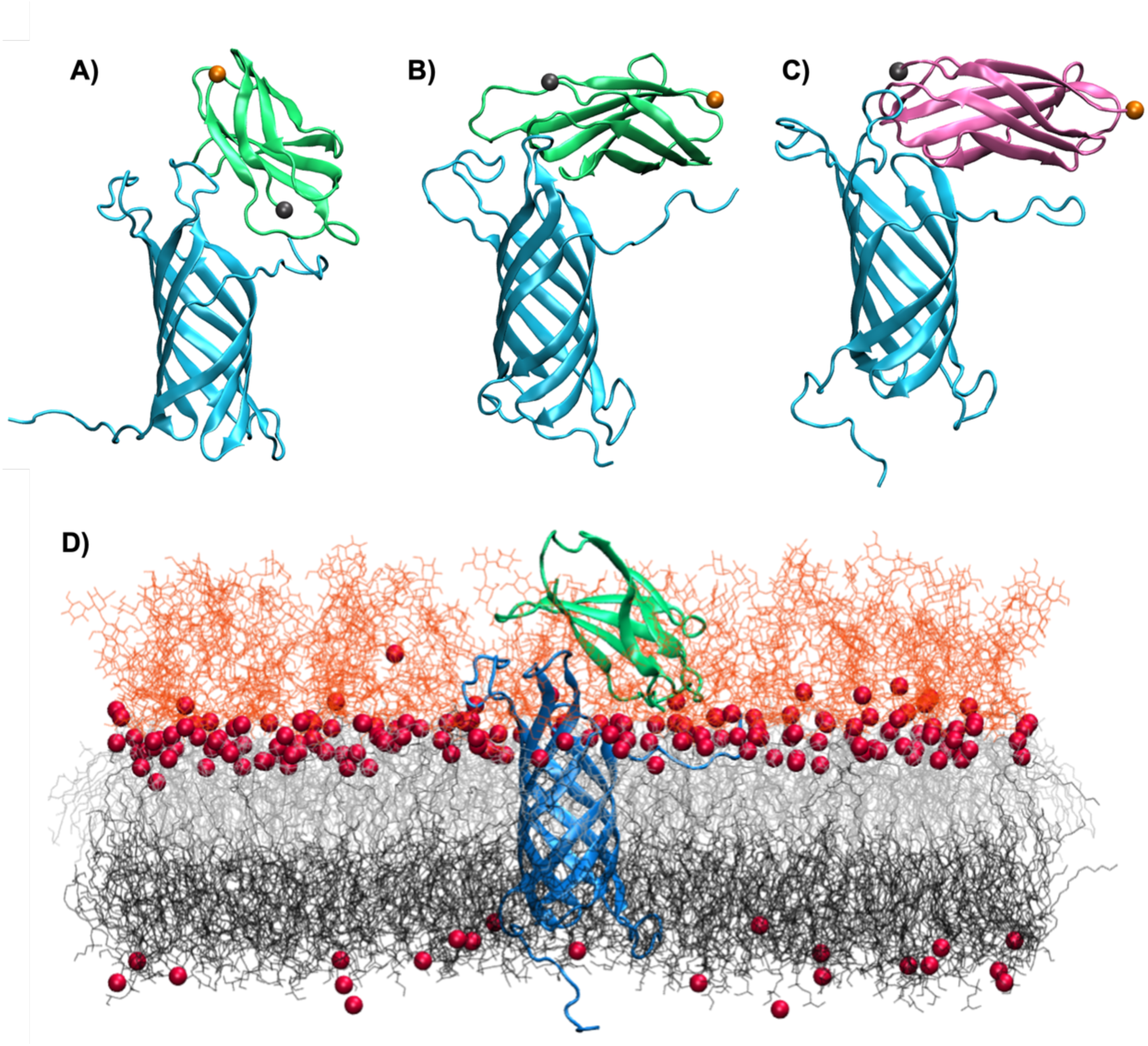
Representative snapshots issued from the MD simulations of the complex between OmpA (in blue) and a unit of FN (green or pink), A) and B) represent simulations performed for two different poses of the RGD-containing unit (RGD_1 and RGD_2), while C represents the simulation from the unique pose for the non-RGD beads. The alpha carbon of head and tail residues of the FN beads are highlighted in VDW sphere respectively in gray and orange. D) Representation of the protein complex embedded in the bacterial membrane including lipopolysaccharides in orange lines, lipids in black and gray, and Ca^2+^ in red and van der Waal representation.

To analyze the results for each amino acid engaged in the complex association we report the percentage of occurrence of specific electrostatic and hydrogen bonds for the different conformations (Tables S1 to S6). Despite the stability of the complex it appears that the individual interactions are rather non persistent, and variable between the different replica. The highest persistence is observed for the aspartate 42 of OmpA interacting with arginine 367 of FN, which is present 39,7% of the simulation time for the RGD_1 configuration (Table S1). However, this situation masks a more complex and dynamic protein/protein interface as described and detailed in Figure 4. Indeed, it appears that some residues are constantly interacting with residues belonging to the other partner, as shown by the distribution of the minimum distance between the residue and the ones from the other protein. In particular, in RGD_1 structure, K360(FN) and D42 (OmpA) are always bridging the protein. This situation is also evident in RGD_2 involving residues K328 (FN) and E134 (OmpA) even if a more bimodal distribution is observed for the latter. The non-RGD conformation follows the same pattern presenting two residues in the FN unit always being in interaction with OmpA (R262 and R238) while E135 (OmpA) is always bridging the FN.

**Figure 4.**
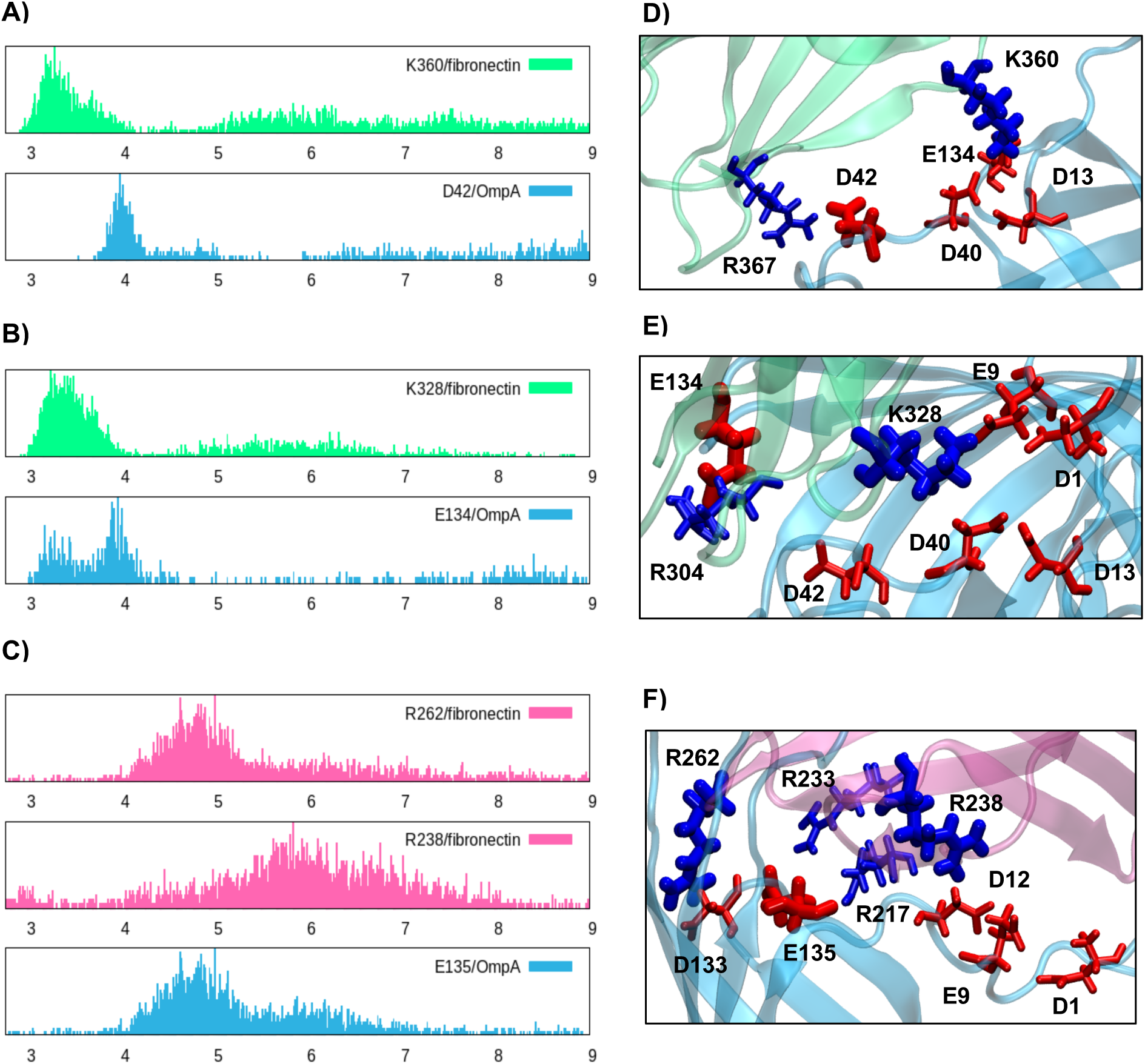
Distribution of the minimum distances between the labeled residue and any residue from the other protein having an opposite charge for A) RGD_1, B) RGD_2, C) non-RGD. E), F) and G) show the ensemble of residues involved in the interaction.

In Figure 4D-F we also identify the main residues involved in the interactions sketched previously. It has been observed that residues are located in close spatial proximity and define an interacting region that stabilizes the contact between the two proteins. As shown from the time series of the relative distances, reported in SI, we may also see that the interaction is dynamic with one residue constantly exchanging partner and exploring the contact interface. This situation is particularly stunning in the case of RGD_2 since K328 is dynamically interacting with 8 other residues. Therefore, we believe that the interaction of OmpA with FN is also a nice example of a dynamic protein/protein interface leading to a large stabilization.

The presence of more important persistent interactions with the non-RGD unit could point to its predominance, yet care should be taken without resorting to the determination of binding free energies. This occurrence could also be biologically significant, since usually RGD is involved in mediating other interactions in the extracellular matrix, and thus could be less available for shielding OmpA.

The previous analysis has strongly confirmed that membrane embedded OmpA is able to persistently interact with one unit of FN through a dynamic interface also leading to polymorphic conformations. However, given that FN is a fibrillar protein composed of multiple FN beads, we have investigated the possibility of making protein/protein complexes by docking two units of FN to OmpA. To avoid reproducing the single unit confirmations, we have specifically defined the contact region at the interface between the two beads. As shown in SI we obtained three independent docking poses. However, at the conlusion of the MD trajectories, only one of them remained viable. Its final structure is displayed in Figure 5. Indeed, in two cases over three the MD became unstable and FN either strongly perturbed the membrane, leading to its unphysical internalization, or destroyed the β-barrel structure of OmpA, also leading to transmembrane water leakage. Obviously, these conformations have been discarded from any further analysis. The stable structure in Figure 5 shares considerable resemblance with the structure of one FN unit interacting with OmpA. FN interacts mainly with the OmpA arms and resides in the polar head and lipopolysaccharide region. As seen in Figure 5A two arms of OmpA interacts with one FN bead, while the third is in contact with the second one. As shown in Figures 5B and 5C the RMSD and RMSF are stable and do not show significant variations compared to the isolated systems.

**Figure 5.**
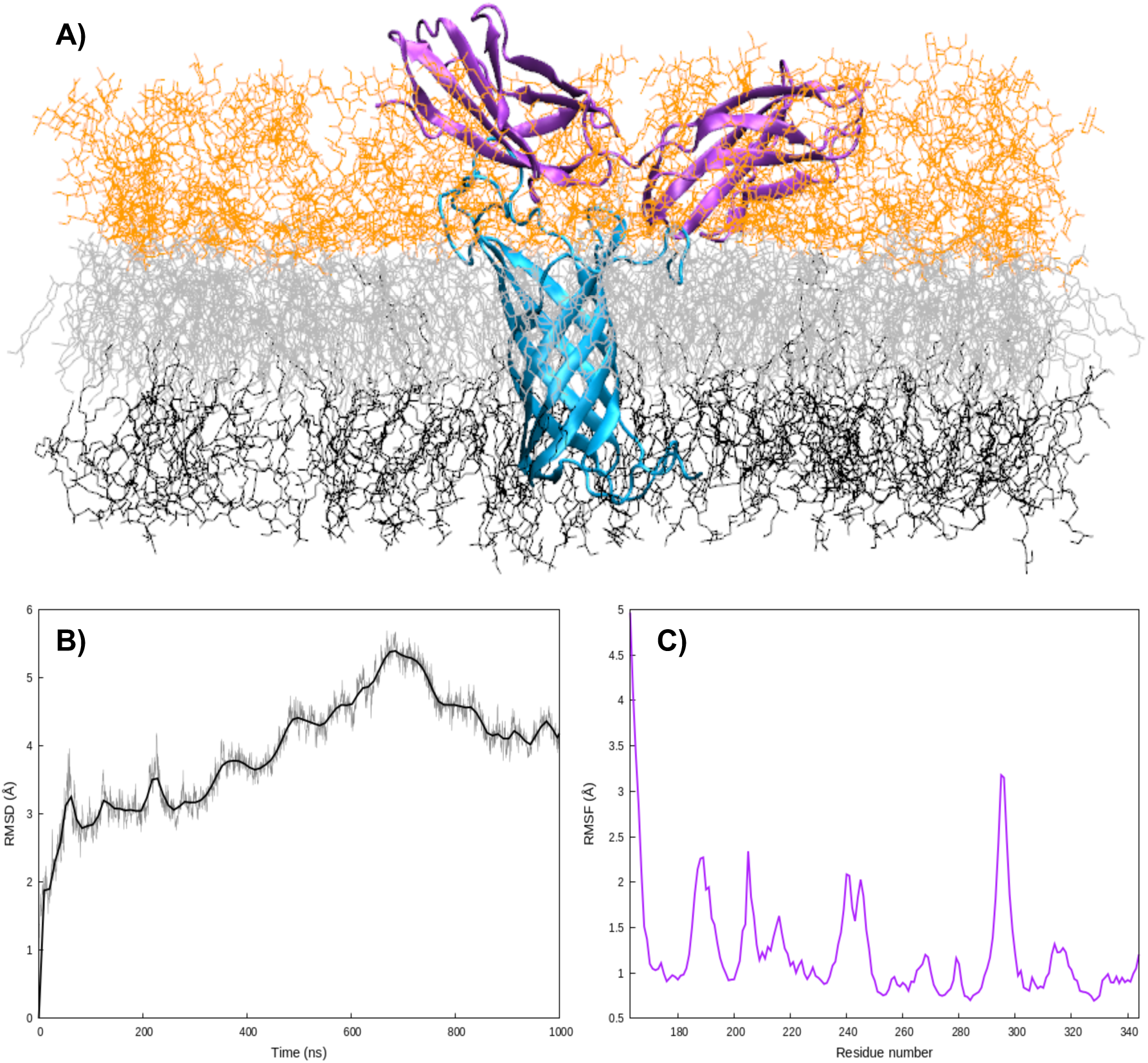
A) Stable conformation for two FN units interacting with membrane embedded OmpA obtained after MD simulation. B) RMSD of the whole complex and C) RMSF of FN.

As concerns the residue-base analysis of the involved amino-acids we may see in Figure 6 that the interface is now more rigid and is mainly mediated by salt bridges between differently charged residues. In particular, OmpA E135 and E134 interacts strongly with FN residues K183 and R262. The second FN unit is instead locked by persistent interactions involving OmpA D12 and D13 with K134 of FN. This pattern is also particularly interesting since it involves residues which have been already pinpointed as crucial to modulate the interaction between FN and OmpA.

**Figure 6.**
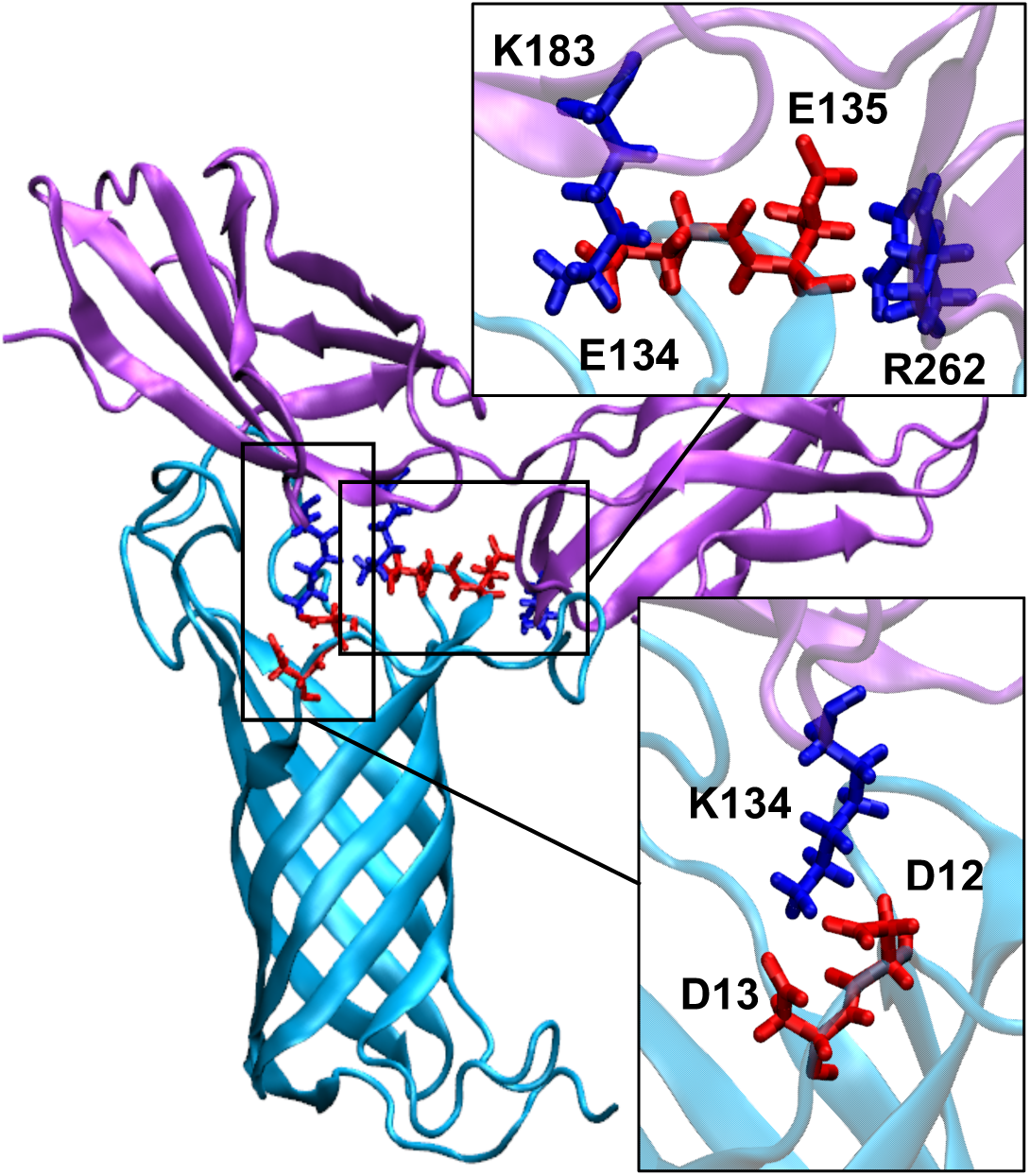
Main electrostatic interactions between OmpA and two FN units as obtained from the MD simulations.

To better analyze the interplay between the one- and two-units interaction patterns we also plot in Figure 7 the superposition of representative MD structures for the two cases. Strikingly, it appears that the two-units complex could be potentially built by the sliding of the non-RGD conformation. More importantly, from the inlay zoom of Figure 7 we may see that OmpA E135 may interact with R262 form both FN units, thus further stabilizing the interface. Note also that this interaction is highly persistent all along the MD simulation.

**Figure 7.**
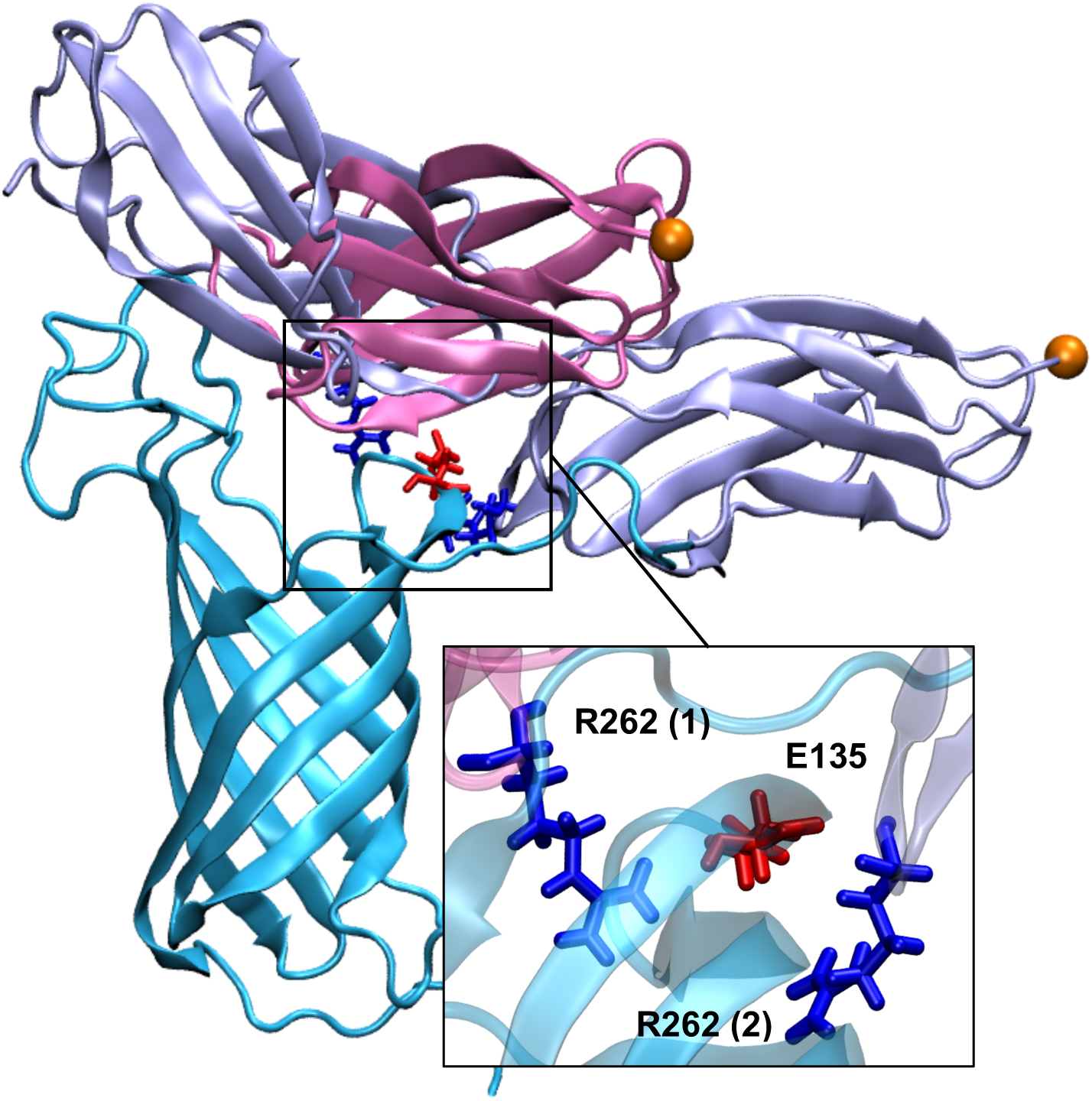
Superposition of OmpA (cyan)/FN complexes structures involving one (pink) and two FN units (ice-blue). The alpha carbon of the FN tail is shown in orange van der Waals. Residues involved in common interactions are shown in licorice representation with blue or red colors for positively or negatively charged, respectively.

From the ensemble of these results it appears that FN may interact with OmpA, mainly involving one unit locked by a dynamic protein/protein interface. However, this situation could also coexist, and probably be in equilibrium with, an interaction involving two FN units which can be accessed by sliding the non-RGD conformation. Even if the interface is complex, it always involves spatially close amino acid, which could, thus, offer attractive drug design hotspot to weaken the OmpA/FN complex and reinstate immune response.

## CONCLUSIONS

The Gramm negative *A. baumannii* is an infectious agent which is raising increasing concern due to its involvement in nosocomial infection and the development of antibiotic resistance. In this contribution, we employed long-range MD simulations to ascertain, for the first time, the structure of the complex between its OmpA external membrane protein and human FN. The formation of this protein/protein complex is meant to shield the bacterium from the immune system recognition, therefore increasing its virulence and resistance. Our findings demonstrate that the complex may mainly involve the interaction with one FN unit, even if this conformation can coexistence with a two-units structure obtained by partially sliding FN on top of OmpA. The protein/protein structure is stabilized by a complex interface characterized by a dynamic interaction network, mainly constituted of salt bridges and hydrogen bonds. We have also identified charged OmpA residues which play a crucial role to mediate the interaction and are conserved in all the conformations. These residues are likely to serve as significant hot-spots for the development of drug-design strategies involving either small molecules, or peptides and aimed at preventing the formation of the complex to reinstate immune response. On the other hand, the RGD domain, which mediated interaction between FN and integrin, does not appear to be involved in the adhesion to the bacterial protein.

Finally, we have also confirmed that OmpA acts as a porin, establishing a continuous water channel through its β-barrel structure, and hence a communication between the periplasm and the extracellular medium. Therefore, OmpA could also be related to antibiotic resistance, allowing the extrusion of drugs. In this respect we plan, in the following, to assess the permeability of OmpA to known antibiotics, such as chloramphenicol through suitable enhanced sampling techniques.

## ASSOCIATED CONTENT

Main docking poses between OmpA and one- or two-FN beads, Detailed tables for the persistence of the interactions in the protein/protein complexes. Time series of the main distances between the interacting residues. Unphysical poses leading to membrane disruption. The following files are available free of charge. (PDF).

## AUTHOR INFORMATION

### Author Contributions

The manuscript was written through contributions of all authors. All authors have given approval to the final version of the manuscript.

## ACKNOWLEDGMENT

L.V. and F.B. kindly acknowledge the financial support of the French ANR under the PIRATE (ANR-21-CE45-0014) project. The authors thank GENCI and Explor computing centers and the Platform P3MB for computational resources. The authors thanks ANR and CGI for their financial support of this work through Labex SEAM ANR 11 LABEX 086, ANR 11 IDEX 05 02 and PIRATE. The support of the IdEx "Université Paris 2019" ANR-18-IDEX-0001.

