## Supplementary Information for "*Acinetobacter baumannii*’s Great Escape: How a Membrane Protein Interacts with Fibronectin to Evade Immunity"

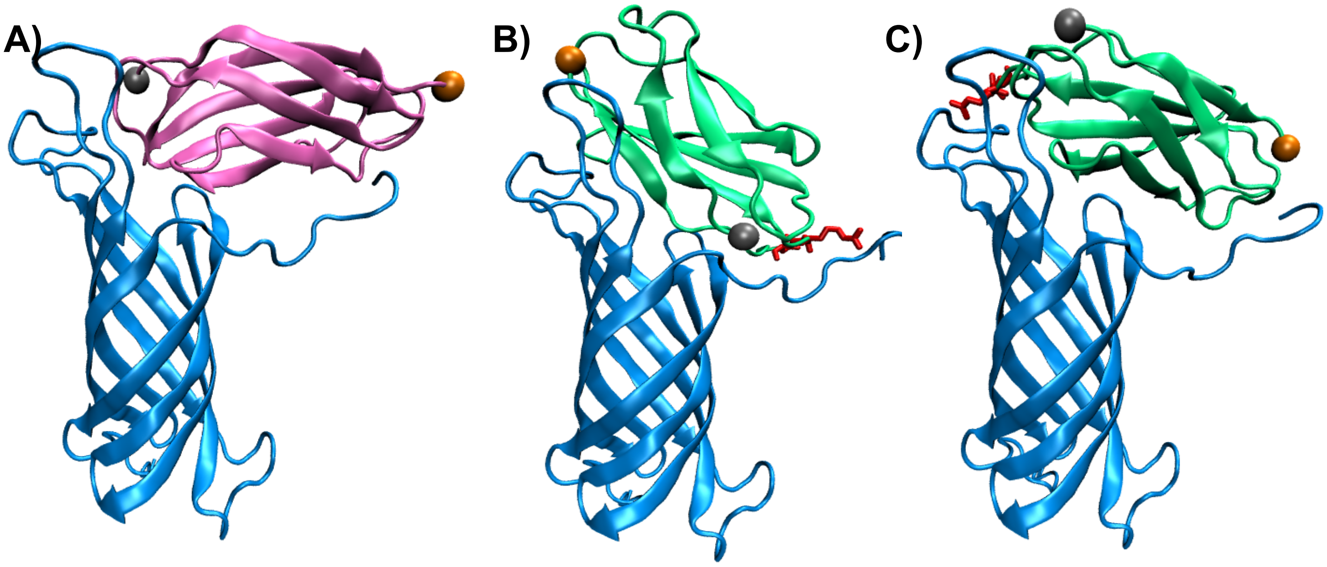


**Figure S1.** Poses of FN/OmpA complex obtained after docking. A) nonRGD (pink), B) RGD_1 (green), C) RGD_2 (green). OmpA is always represented in blue.

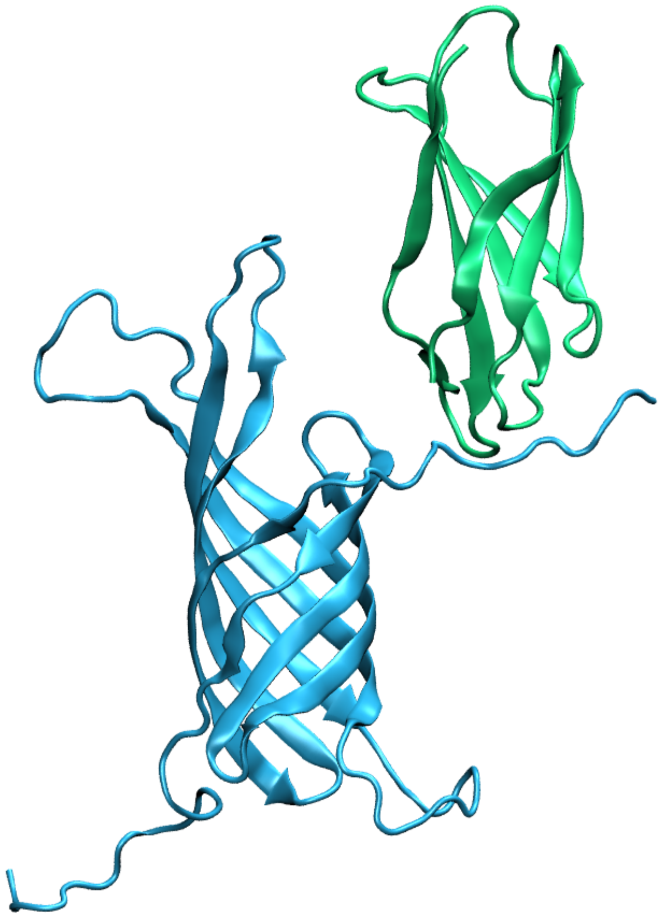

**Figure S2.** Structure of the unphysical binding between OmpA and RGD_1. This structure has been discarded from further analysis since the structure is not compatible with an extended FN unit. OmpA is represented in blue and RGD_1 in green.

| RGD_1 | R1 | R2 | R3 | R4 | Total |
|---|---|---|---|---|---|
| **D42 - R367** | 0.81 | 99.01 | 0.1 | 20.1 | **39.7** |
| **D1 - R352** | 2.15 | 0.04 | 17.3 | 74.6 | **30.6** |
| **E9 - R280** | 0 | 0.08 | 0.2 | 91.4 | **30.6** |
| **E134 - K360** | 0.22 | 0 | 22.0 | 36.8 | **19.6** |
| **D40 - K360** | 6.15 | 6.07 | 40.4 | 4.1 | **16.9** |
| **D13 - K360** | 2.52 | 5.59 | 3.0 | 23.0 | **10.5** |
| **D12 - R280** | 1.15 | 1.47 | 0.0 | 22.9 | **8.1** |

**Table S1.** Percentage of presence of electrostatic interactions between OmpA and RGD_1, over the 1µs of simulation time. Gray cases report the values for the unphysical replica not considered (see Figure S2). The color of cases depending on the persistence of the interaction, blue: <7%; green: 7%≤x<20%; yellow: 20%≤x<40%; orange: 40%≤x<60% and red: ≥ 60%.

| RGD_2 | R1 | R2 | R3 | R4 | Total |
|---|---|---|---|---|---|
| **E134 - K328** | 17.7 | 13.7 | 88.7 | 31.8 | **38.0** |
| **E134 - R304** | 17.8 | 0.0 | 0.0 | 72.6 | **22.6** |
| **D1 - K328** | 0.0 | 0.0 | 0.0 | 80.7 | **20.2** |
| **D40 - K328** | 4.0 | 17.3 | 0.0 | 42.0 | **15.8** |
| **E101 - K328** | 0.0 | 49.5 | 2.7 | 0.4 | **13.2** |
| **E9 - K328** | 40.7 | 0.0 | 0.1 | 5.7 | **11.6** |
| **E101 - R352** | 0.0 | 0.0 | 0.0 | 44.2 | **11.0** |
| **D42 - K328** | 0.1 | 21.2 | 0.0 | 22.6 | **11.0** |
| **D13 - K328** | 5.2 | 1.2 | 0.7 | 28.5 | **8.9** |

**Table S2.** Percentage of presence of electrostatic interactions between OmpA and RGD_2, over the 1μs of simulation time. The color of cases depending on the persistence of the interaction, blue: <7%; green: 7%≤x<20%; yellow: 20%≤x<40%; orange: 40%≤x<60% and red: ≥ 60%.

| nonRGD | R1 | R2 | R3 | R4 | Total |
|---|---|---|---|---|---|
| **E135 - R262** | 51.4 | 7.0 | 67.2 | 5.5 | **32.8** |
| **E135 - R233** | 27.8 | 0.6 | 12.2 | 81.3 | **30.5** |
| **D133 - R262** | 51.2 | 8.1 | 45.8 | 6.4 | **27.9** |
| **E9 - R238** | 19.2 | 6.1 | 25.0 | 44.5 | **23.7** |
| **D12 - R238** | 24.0 | 2.0 | 23.9 | 4.5 | **13.6** |
| **D1 - R228** | 0.0 | 0.0 | 0.1 | 41.2 | **10.3** |
| **E135 - R217** | 19.0 | 13.8 | 0.6 | 0.2 | **8.4** |

**Table S3.** Percentage of presence of electrostatic interactions between OmpA and nonRGD, over the 1μs of simulation time. The color of cases depending on the persistence of the interaction, blue: <7%; green: 7%≤x<20%; yellow: 20%≤x<40%; orange: 40%≤x<60% and red: ≥ 60%.

| RGD_1 | R1 | R2 | R3 | R4 | Total |
|---|---|---|---|---|---|
| **E9 - D277** | 37.7 | 40.3 | 28.76 | 62.84 | **43.98** |
| **E9 - R280** | 0 | 0 | 0 | 104.84 | **34.95** |
| **D42 - R367** | 0.3 | 58.2 | 0 | 22.56 | **26.93** |
| **D40 - S363** | 0.0 | 65.5 | 0 | 0 | **21.82** |
| **Y42 - N365** | 0.1 | 46.5 | 0 | 0 | **15.50** |
| **Y130 - S355** | 0.0 | 37.7 | 0 | 0 | **12.55** |

**Table S4.** Percentage of presence of hydrogen bonds between OmpA and RGD_1, over the 1μs of simulation time. Gray cases report the values for the unphysical replica not considered (see Figure S2). The color of cases depending on the persistence of the interaction, blue: <7%; green: 7%≤x<20%; yellow: 20%≤x<40%; orange: 40%≤x<60% and red: ≥ 60%.

| RGD_2 | R1 | R2 | R3 | R4 | Total |
|---|---|---|---|---|---|
| **E134 - R304** | 6.0 | 0.0 | 0.0 | 56.5 | **15.6** |
| **E134 - K328** | 2.2 | 0.0 | 39.8 | 0.0 | **10.5** |
| **D1 - T288** | 34.6 | 0.0 | 0.0 | 0.0 | **8.7** |
| **D1 - K328** | 0.0 | 0.0 | 0.0 | 32.9 | **8.2** |

**Table S5.** Percentage of presence of hydrogen bonds between OmpA and RGD_2, over the 1μs of simulation time. The color of cases depending on the persistence of the interaction, blue: <7%; green: 7%≤x<20%; yellow: 20%≤x<40%; orange: 40%≤x<60% and red: ≥ 60%.

| nonRGD | R1 | R2 | R3 | R4 | Total |
|---|---|---|---|---|---|
| **E134 - T214** | 28.7 | 9.2 | 35.9 | 9.8 | **20.9** |
| **E134 - R233** | 9.4 | 0.0 | 5.2 | 48.6 | **15.8** |
| **E135 - R233** | 15.9 | 33.2 | 7.7 | 4.5 | **15.3** |
| **D1 - D232** | 0.1 | 0.0 | 44.4 | 12.6 | **14.3** |
| **E134 - G261** | 20.5 | 5.6 | 1.8 | 27.0 | **13.7** |
| **G2 - D232** | 0.0 | 0.0 | 41.8 | 5.2 | **11.8** |
| **T5 - T242** | 0.1 | 0.0 | 40.1 | 6.3 | **11.6** |
| **D133 - R262** | 26.9 | 2.0 | 14.3 | 2.7 | **11.5** |
| **E135 - R217** | 4.1 | 37.4 | 0.0 | 0.0 | **10.4** |
| **E101 - T212** | 0.2 | 0.0 | 32.4 | 6.3 | **9.7** |
| **Y50 - N260** | 24.7 | 3.3 | 4.7 | 0.0 | **8.2** |

**Table S6.** Percentage of presence of hydrogen bonds between OmpA and nonRGD, over the 1μs of simulation time. The color of cases depending on the persistence of the interaction, blue: <7%; green: 7%≤x<20%; yellow: 20%≤x<40%; orange: 40%≤x<60% and red: ≥ 60%.

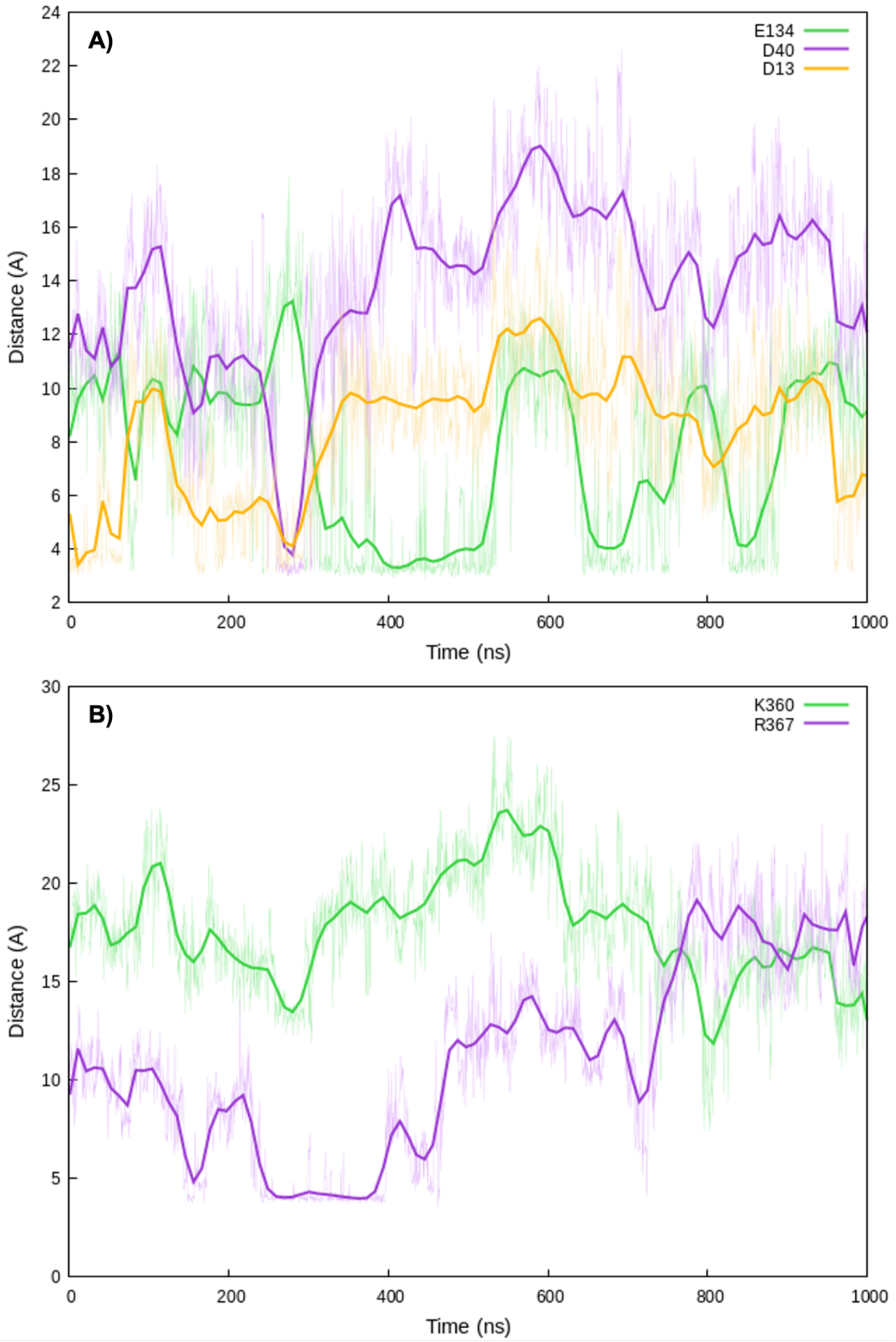


**Figure S3.** Timeseries of the distance between the residue K360 A)  and residue D42 B) and all the closest residues for the RGD_1 complex.

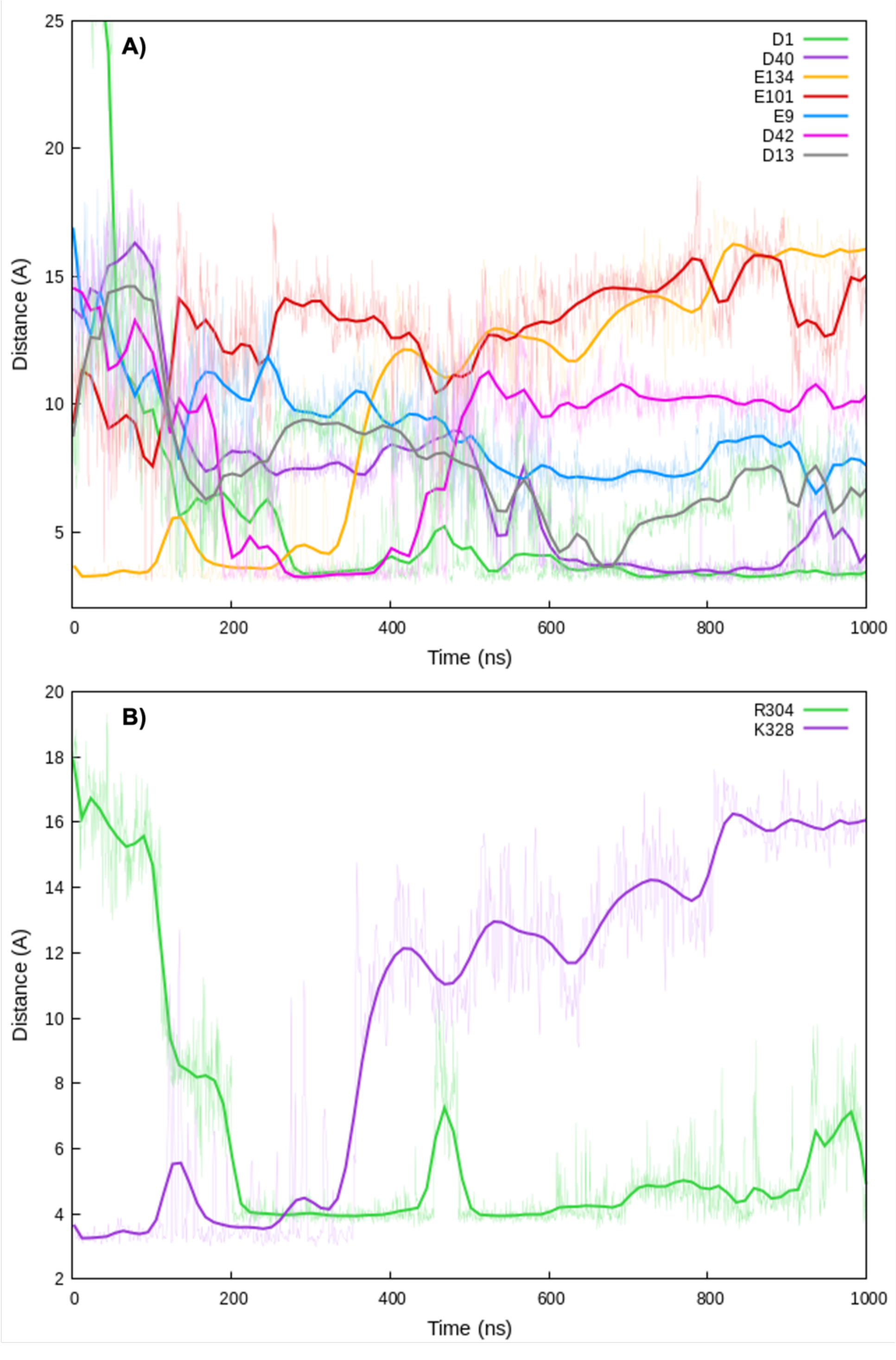


**Figure S4.** Timeseries of the distance between the residue K328 A) and residue E134 B) and all the closest residues for the RGD_2 complex.

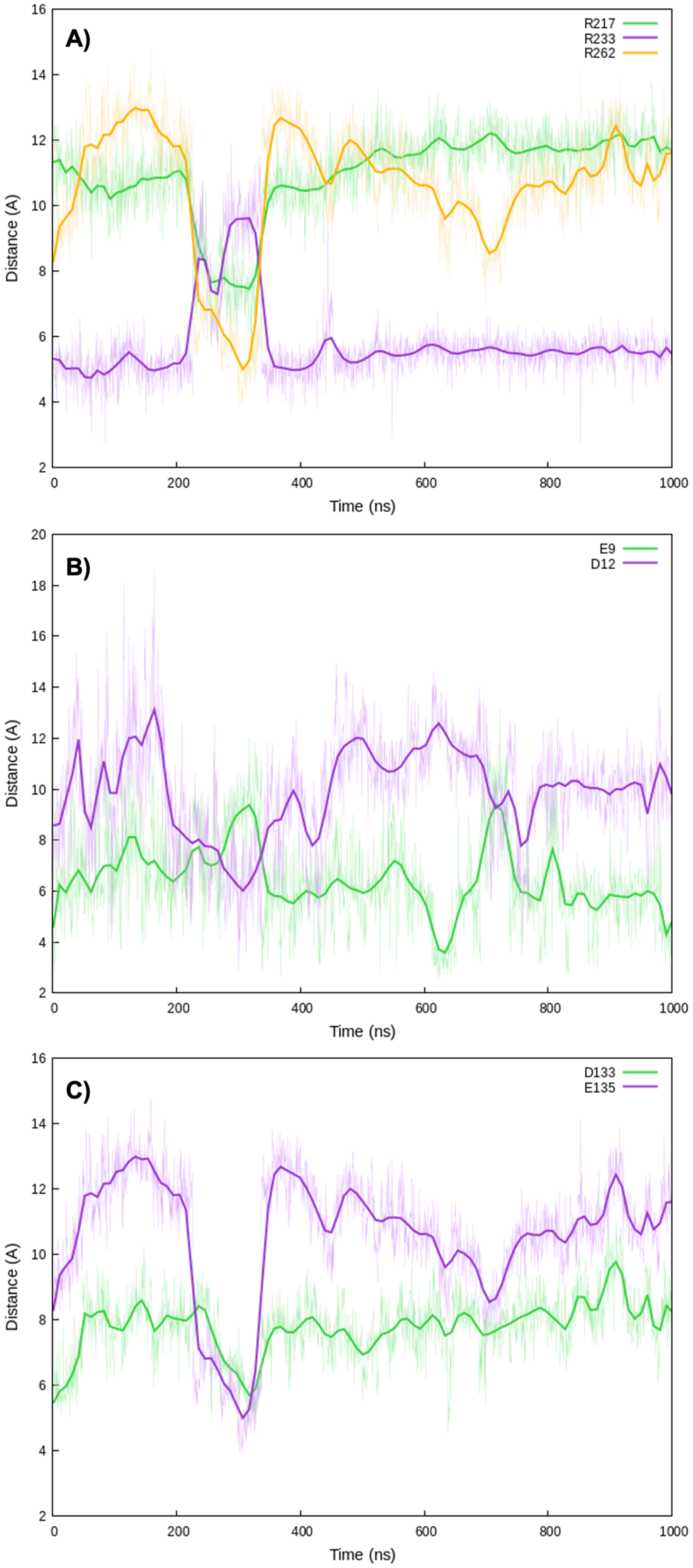


**Figure S5.** Timeseries of the distance between the residue R262 A), residue R262 B), and residue R238 C) and all the closest residues for the non-RGD complex.

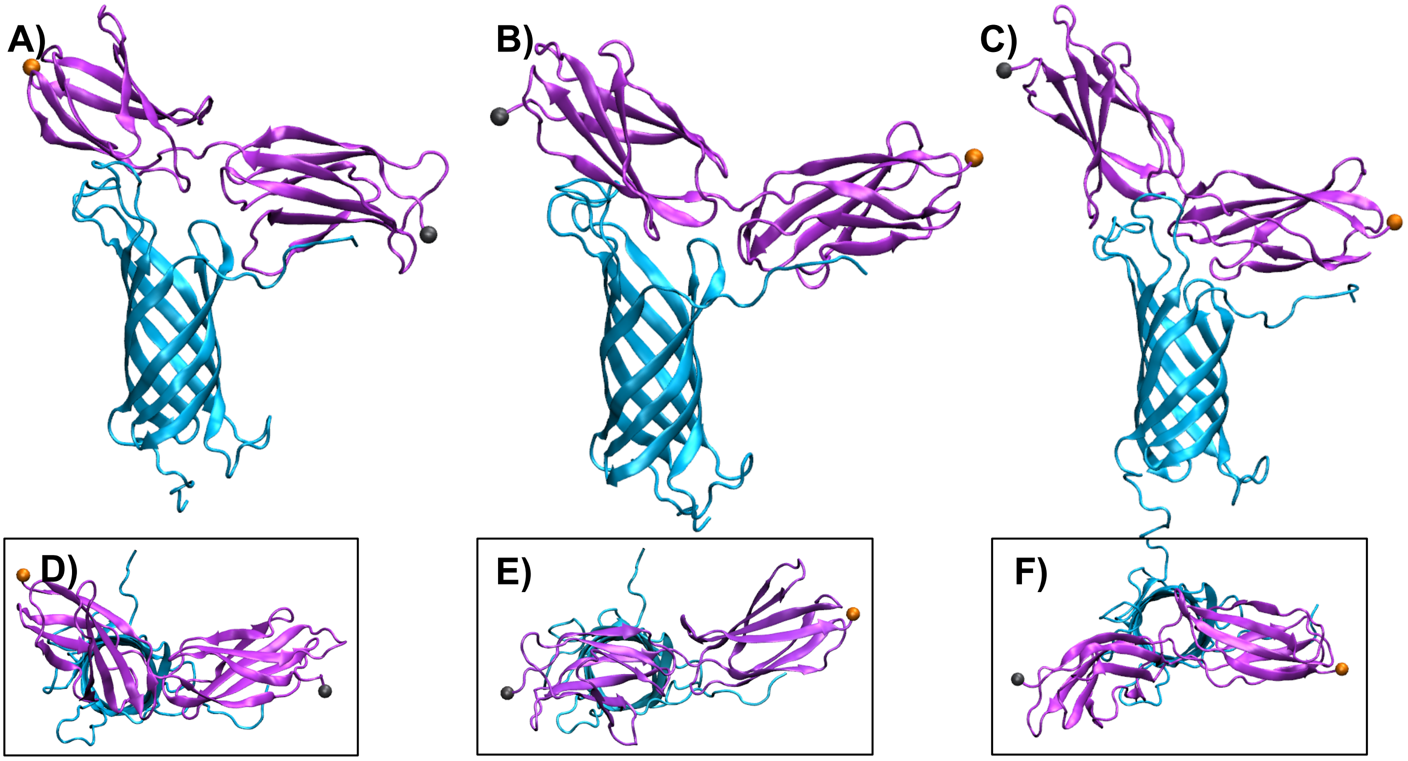


**Figure S6.** Docking poses for the complex between OmpA (blue) and two non-RGD beads of FN (purple). D), E) and F) are the top view of A), B) and C) respectively.

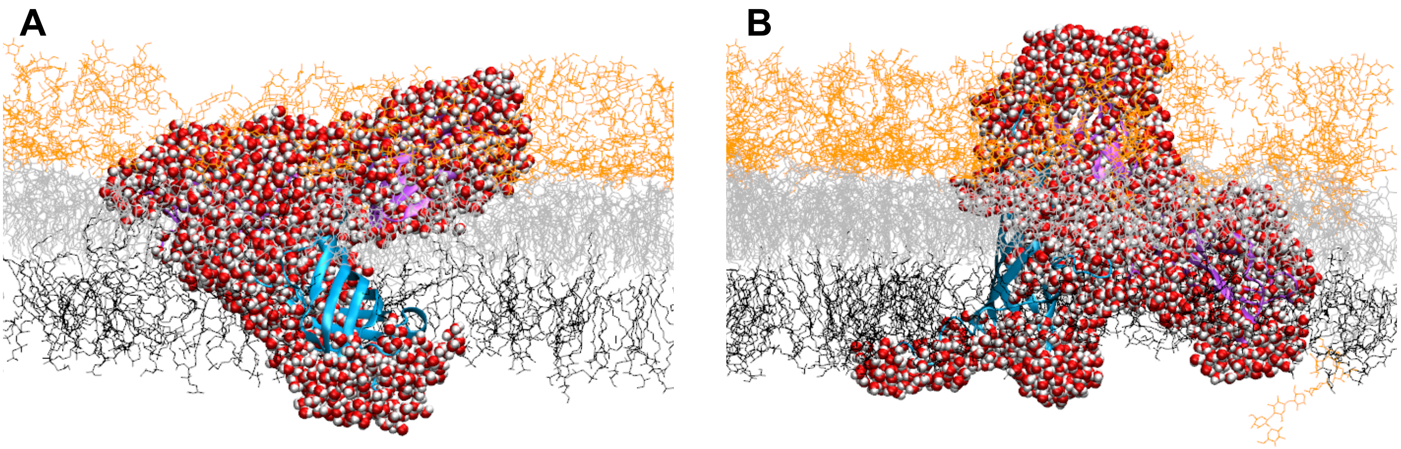


**Figure S7.** Examples of rejected structures due to membrane disruption and permeabilization. Lipid in gray and black, lipopolysaccharides in orange, OmpA in blue, two-unit nonRGD of fibronectin in purple. Red and white: water molecules.
